# Modeling differences in neurodevelopmental maturity of the reading network using support vector regression on functional connectivity data

**DOI:** 10.1101/2024.06.10.597945

**Authors:** Oliver H.M. Lasnick, Jie Luo, Brianna Kinnie, Shaan Kamal, Spencer Low, Natasza Marrouch, Fumiko Hoeft

## Abstract

The construction of growth charts trained to predict age or developmental deviation (the ‘brain-age index’) based on structural/functional properties of the brain may be informative of children’s neurodevelopmental trajectories. When applied to both typically and atypically developing populations, results may indicate that a particular condition is associated with atypical maturation of certain brain networks. Here, we focus on the relationship between reading disorder (RD) and maturation of functional connectivity (FC) patterns in the prototypical reading/language network using a cross-sectional sample of N = 742 participants aged 6-21 years. A support vector regression model is trained to predict chronological age from FC data derived from a whole-brain model as well as multiple ‘reduced’ models, which are trained on FC data generated from a successively smaller number of regions in the brain’s reading network. We hypothesized that the trained models would show systematic underestimation of brain network maturity for poor readers, particularly for the models trained with reading/language regions. Comparisons of the different models’ predictions revealed that while the whole-brain model outperforms the others in terms of overall prediction accuracy, all models successfully predicted brain maturity, including the one trained with the smallest amount of FC data. In addition, all models showed that reading ability affected the ‘brain-age’ gap, with poor readers’ ages being underestimated and advanced readers’ ages being overestimated. Exploratory results demonstrated that the most important regions and connections for prediction were derived from the default mode and frontoparietal control networks.

**Glossary:** Developmental dyslexia / reading disorder (RD): A specific learning disorder affecting reading ability in the absence of any other explanatory condition such as intellectual disability or visual impairment

Support vector regression (SVR): A supervised machine learning technique which predicts continuous outcomes (such as chronological age) rather than classifying each observation; finds the best-fit function within a defined error margin

Principal component analysis (PCA): A dimensionality reduction technique that transforms a high-dimensional dataset with many features per observation into a reduced set of ‘principal components’ for each observation; each component is a linear combination of several original (correlated) features, and the final set of components are all orthogonal (uncorrelated) to one another

Brain-age index: A numerical index quantifying deviation from the brain’s typical developmental trajectory for a single individual; may be based on a variety of morphometric or functional properties of the brain, resulting in different estimates for the same participant depending on the imaging modality used

Brain-age gap (BAG): The difference, given in units of time, between a participant’s true chronological age and a predictive model’s estimated age for that participant based on brain data (Actual – Predicted); may be used as a brain-age index

*Highlights:*

- A machine learning model trained on functional data predicted participants’ ages
- The model showed variability in age prediction accuracy based on reading skills
- The model highly weighted data from frontoparietal and default mode regions
- Neural markers of reading and language are diffusely represented in the brain

## 1. Introduction

The development of ‘brain growth charts’ derived from neuroimaging data allows neuroscientists to examine properties of the brain, such as regional gray/white matter density or functional magnetic resonance imaging (fMRI) activity (an indirect measure of metabolic activity; Ogawa et al., 1993) and observe how these metrics change across time (Giorgio et al., 2008; Lemaitre et al., 2012; Bray et al., 2015). Tracking atypical trajectories of brain development is crucial for understanding the progression of neurodevelopmental disorders (Wang et al., 2021). Researchers have previously tracked the maturation of brain structure and function from childhood to adulthood (Fair et al., 2007; Fair et al., 2008; Dong et al., 2020). Regions in the default mode network (DMN), a network that is most active during wakeful rest and posited to play diverse roles in social cognition, memory, and multimodal integration (Smallwood et al., 2021), shift from being sparsely interconnected to densely interconnected with increasing age (Buckner et al., 2008; Fair et al., 2008). In general, short-range connections tend to be negatively correlated with age (decrease in strength from childhood to adulthood), while long-range connections grow stronger with age (Fair et al., 2007).

More recent advances include machine learning (ML)-based growth curves. Dosenbach et al. (2010) used support vector regression (SVR) to train a model to predict brain age/maturity from functional connectivity (FC) data. SVR is a supervised learning algorithm that predicts continuous outcomes by finding the best-fit function within a defined error margin (Drucker et al., 1997). Such a brain-age index can be used to track developmental deviation across time and within clinical populations, such as those with ADHD (Kessler et al., 2016). Such models could reveal a key source of neurodevelopmental deviation in atypically-developing populations. If a child has altered early brain development, their estimated brain age could differ substantially from their chronological age, which is reflected as a larger brain age gap (BAG, An et al., 2025). For example, two meta-analyses concluded that children who experience trauma or adverse childhood events (ACEs) show higher DMN activity during a broad swathe of tasks (emotional/social processing, memory, and self-referential thought) than control children (Ireton et al., 2024), in addition to signs of accelerated neurobiological aging (cortical thinning) in the prefrontal/frontoparietal cortex and the DMN (Colich et al., 2020).

One area where the brain-age index remains underexplored is in developmental dyslexia, or specific reading disorder (RD) (Peterson & Pennington, 2015; Swagerman et al., 2017). RD is characterized by difficulties in reading that cannot be explained by intellectual disability or sensory perception deficits. In this context, deviations in brain age for RDs compared to their same-age, typically developing peers may indicate atypical maturation in neural systems essential for reading. A younger estimated brain age, for example, could suggest the brain has not reached the biological stage needed for fluent reading, potentially explaining the persistent difficulties seen in children with RD despite adequate instruction. Importantly, brain age estimates can be derived from either whole-brain data (traditional approach) or from distinct sets of brain regions. Isolated underdevelopment (larger BAG) of reading-related brain regions that does not extend to other brain regions could explain the specific reading and language deficits observed in RD. Reading recruits widely distributed left-hemisphere ventral occipitotemporal, inferior frontal, and posterior parietal regions in both children and adults (Martin et al., 2015; Bailey et al., 2018). Those with RD have reduced functional activity in inferior parietal cortex, the superior temporal gyrus, and the inferior frontal gyrus; reduced gray matter volume in the superior temporal gyrus, temporoparietal cortex, middle frontal gyrus, and superior occipital gyrus; and reduced white matter in bilateral parieto-occipital regions (Maisog et al., 2008; Xia et al., 2016; Linkersdörfer et al., 2015). Many of these regions are part of the frontoparietal attention network and the DMN (Bailey et al., 2018); models trained to predict age from data in these regions may also capture brain features associated with reading ability.

To the best of our knowledge, no studies to date have constructed brain-based growth charts for age prediction, allowing for comparison of trajectories among high-, mid-, and low-level readers. Such a model could reveal population-level deviations in children who later meet diagnostic criteria for RD. Beyond methodological value, brain-age estimation offers a theoretical framework to distinguish whether RD reflects a maturational lag that is brain-wide or isolated to specific reading-related regions. Such markers may enable earlier detection of RD risk than behavioral screening alone.

Our first goal was to construct a model that successfully predicts true age in a cross-sectional sample of children with a wide range of reading ability. Our second goal was to perform an exploratory analysis of which brain regions and connections drive model performance. To achieve these goals, we train a linear SVR model on FC data from the whole brain and investigate the relative importance of each functional connection based on model coefficients. Our third goal was to determine whether such a model is able to distinguish the developmental trajectories of exceptional, typical, and impaired readers. To achieve our third goal, we systematically vary training features, generating multiple models trained with data from either whole-brain or narrow (reading-specific) brain regions, and investigate whether model estimates of brain age and/or the BAG differ based on participants’ reading ability. Our sample includes precocious readers (children with unexpectedly advanced reading skills for their age or IQ; Ostrolenk et al., 2017), with the expectation that this sample might also show altered patterns of development in comparison to the typical-range and impaired readers (see Methods).

We hypothesize that the models will be biased in their age estimates for both the impaired readers and precocious readers, reflecting a developmental delay for the former and a developmental acceleration for the latter. We also expect that the whole-brain model will better predict age *in general* (collapsed across reading ability) compared to the reading-network trained models, due to being trained on more data. However, we further hypothesize that group bias in brain age predictions will be moderated by model type (whole-brain vs. reduced): we expect there to be less bias in the whole-brain model (with similar prediction accuracy across reading groups) compared to the reduced models, for which there will be greater prediction errors in the impaired and precocious readers compared to typical readers (i.e., developmental differences in brain connectivity associated with reading ability will be more overtly expressed in the brain’s reading network).

## 2. Materials and Methods

### 2.1. Participants

All data is from the Child Mind Institute (CMI) Healthy Brain Network (HBN) Project, which collects neuroimaging and phenotypic data from a diverse sample of participants (Alexander et al., 2017). Neuroimaging data is publicly available for download on the CMI HBN website, while phenotypic data was made available for download via the Longitudinal Online Research and Imaging System (LORIS) upon completion of a Data Usage Agreement (DUA).

#### 2.1.1. Inclusion criteria

We included participants who were ≥ 6 years of age at the time of data collection (6-21 years). The Test of Word Reading Efficiency (TOWRE), designed for participants aged 6-24 years, was used for group classification criteria (Torgesen et al., 2012). TOWRE involves two core subtests: Phonemic Decoding Efficiency (PDE) and Sight Word Efficiency (SWE). The researcher presents a participant with a list of either real words or pseudowords, which the participant is asked to read aloud one-by-one as quickly as possible within the allotted time. Real word reading is thought to index sight word reading ability, while pseudoword reading is thought to index phonemic decoding skills. The composite Total Word Reading Efficiency (TWRE) index is derived from these two subtests (Figure 1). A participant is considered a poor reader (PR) if TWRE ≤ 90, a typical reader (TR) if 91 ≤ TWRE ≤ 109, and an exceptional reader (ER) if TWRE ≥ 110.

**Figure 1.**
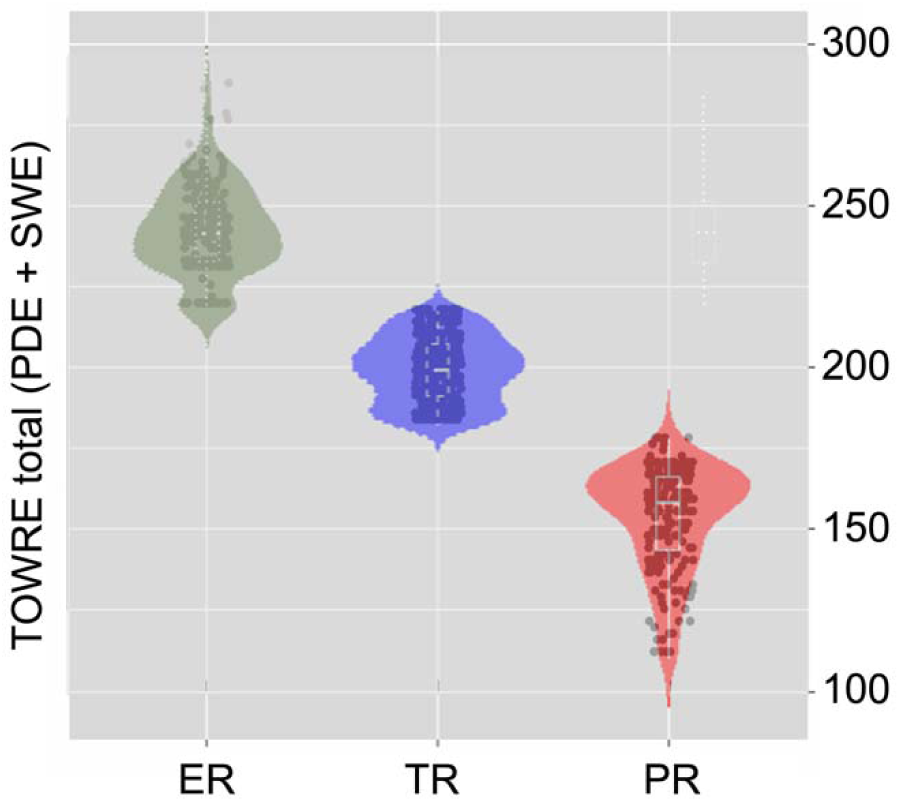
TOWRE total scores across each group. Individual dots represent a single participant; violin plots are used to show the distribution of scores within each group.

#### 2.1.2. Comorbidities

We do not exclude participants based on certain comorbid clinician diagnoses. This was done to avoid the emergent population-level bias in traditional studies which disallow this kind of natural variation. RD has high comorbidity with attention deficit hyperactivity disorder (ADHD) and other specific learning disorders, such as writing (dysgraphia) and mathematics (dyscalculia), as well as internalizing disorders such as depression and anxiety (Hendren et al., 2018). In addition, while RD has often been defined in terms of unexpectedly poor decoding in the absence of intellectual disability (Rutter & Yule, 1975), associations between RD and IQ scores (both verbal and nonverbal) have been reported at the population level, particularly for verbal IQ (van Bergen et al., 2014).

Table 1 contains all demographic and behavioral data on the participants after scan quality-control criteria had been applied. Those demographic variables with significant differences between groups (as well as gender and handedness) were included as covariates/nuisance regressors for all analyses, except for TOWRE subtest scores, which were used as grouping criteria; and full-scale IQ (FSIQ), which was highly correlated with performance (nonverbal) IQ (PIQ). Covariates were age, gender, handedness, PIQ, and SES.

**Table 1.** Demographic & behavioral data.

|  | **ER**<br>**N = 193** | **TR**<br>**N = 338** | **PR**<br>**N = 211** | **Test statistic** | **Effect size**<br>**($\eta^2$)** |
| --- | --- | --- | --- | --- | --- |
| **Age**<sup>1¶</sup> | 10.2 (2.9)<sub>a</sub> | 10.7 (2.7)<sub>a,b</sub> | 11.2 (3.6)<sub>b</sub> | Welch's F(2,407.616) = 4.88 ($p = .008$)** | .015 |
| **Gender (% female)**<sup>¶</sup> | 29.5 | 36.7 | 34.6 | $\chi^2(2,739) = 2.81$ ($p = .245$) | |
| **Socioeconomic status (SES)**<sup>2¶</sup> | 9.0 (3.0)<sub>a</sub><br>[N=166] | 8.5 (3.1)<sub>b</sub><br>[N=280] | 6.8 (3.7)<sub>c</sub><br>[N=158] | Kruskal-Wallis H(2) = 37.85 ($p < .001$)*** | .060 |
| **Handedness (% right)**<sup>¶</sup> | 91.7 | 86.7 | 91.0 | $\chi^2(2,739) = 4.19$ ($p = .123$) | |
| **FSIQ**<sup>1</sup> | 114.0 (14.4)<sub>a</sub> | 102.6 (12.1)<sub>b</sub> | 90.8 (12.6)<sub>c</sub><br>[N=178] | Welch's F(2,380.667) = 136.66 ($p < .001$)*** | .298 |
| **PIQ (Block design, Scaled)**<sup>¶</sup> | 11.6 (3.1)<sub>a</sub> | 10.3 (3.2)<sub>b</sub> | 8.9 (2.9)<sub>c</sub><br>[N=178] | F(2,706) = 36.39 ($p < .001$)*** | .093 |
| **TOWRE** |  |  |  |  |  |
| **PDE (Scaled)**<sup>1</sup> | 118.0 (8.4)<sub>a</sub> | 98.4 (7.6)<sub>b</sub> | 76.1 (9.3)<sub>c</sub> | Welch's F(2,411.073) = 1139.03 ($p < .001$)*** | .778 |
| **SWE (Scaled)**<sup>1</sup> | 123.8 | 100.7 | 77.5 | Welch's F(2,408.330) = 1269.40 ($p$ | .797 |
|  | (8.6)<sub>a</sub> | (7.8)<sub>b</sub> | (10.0)<sub>c</sub> | <.001)<sup>***</sup> |  |

<sup>1</sup> Levene's test $p < .05$ ; Welch's statistic and Games-Howell correction used.
<sup>2</sup> Kruskal-Wallis nonparametric test reported for ordinal variable; Mann-Whitney U test for pairwise follow-ups.
<sup>¶</sup>Included as covariates for analyses.
Cells with the same subscript indicate non-significant group comparisons at the .05 alpha level. All cells are in the format MEAN (SD) except for gender and handedness, which are given in percentages. IQ was measured using the Wechsler Intelligence Scale for Children – Fifth Edition (WISC-V). Post-hoc tests use Bonferroni correction for multiple comparisons if Levene's test is not significant. Where data is missing, actual N is given in brackets.

### 2.2. Resting fMRI data acquisition

fMRI data was collected during a movie-watching paradigm: the participants viewed a 10-minute-long clip from the film *Despicable Me* while in-scanner. The clip was presented as part of a longer in-scanner protocol and played at the following time stamp: 01:02:09–01:12:09 (HH:MM:SS). The protocol script was written in PsychoPy2 Experiment Builder v1.83.04 (Peirce, 2007). Details of scanner parameters are given in Supplementary File 1. We chose to use this naturalistic movie-watching paradigm rather than traditional resting-state because previous literature suggests that such paradigms may produce data with higher reliability than intrinsic resting-state and even task-based fMRI (Wang et al., 2017; Aliko et al., 2020).

### 2.3. Preprocessing

Data was downloaded from the CMI HBN Project neuroimaging data portal at http://fcon_1000.projects.nitrc.org/indi/cmi_healthy_brain_network/ (DOI: 10.1038/sdata.2017.181). Data preprocessing (intensity non-uniformity correction, skull-stripping, and segmentation for anatomical T1 data; susceptibility distortion correction and co-registration for functional data) and spatial registration of anatomical and functional data were performed using fMRIPrep v1.5.8 (Esteban et al., 2019). For detailed information on the fMRIPrep preprocessing pipeline see Supplementary File 1. Regression of confounds from the BOLD signal was performed using the CONN functional connectivity toolbox (Whitfield-Gabrieli & Nieto-Castanon, 2012). Subcortical, white matter, and cerebrospinal fluid signals derived from tCompCor and aCompCor; framewise displacement (FD); and realignment parameters were regressed from the BOLD signal and high-motion frames were removed (‘scrubbing’, described in Power et al., 2012). A bandpass filter of 0.008-0.09 Hz was then applied to the signal, and linear detrending was applied to correct for time-dependent artifactual signal drift.

### 2.4. Quality control & exclusion criteria

Participants were excluded if (1) their functional data did not include a full run of the movie-watching paradigm (e.g., left the scanner early), or (2) their anatomical T1 scan or functional run was determined to be of insufficient quality. For T1 data, MRIQC was used to accept or reject scans based on quality assessed using a 64-feature vector derived from 14 image quality metrics (Esteban et al., 2017). For functional data, the criteria for exclusion was having more than one-third of frames flagged for excess motion (>0.5mm FD) by fMRIPrep. This left a final sample size of N = 742 participants. A Pearson Chi-squared test found that the rate of exclusion differed between groups: *X***^2^**(2,939) = 9.77 (*p* = .008). Post-hoc tests found that the difference was driven by TRs and PRs: *X***^2^**(1,691) = 9.62 (*p* = .002), with the PRs more likely to be excluded. The difference between TRs/ERs (*X***^2^**(1,653) = 3.21; *p* = .079) and PRs/ERs (*X***^2^**(1,534) = 1.18; *p* = .312), were non-significant.

### 2.5. Regions of interest (ROIs)

This study utilized the Schaefer et al. (2018) 400-ROI seven-network 2mm Montreal Neurological Institute (MNI) template to segment the brain into 400 distinct ROI time series, with networks based on the functional seven-network template described in Yeo et al. (2011). The seven networks are Visual (Vis), Somatomotor (SomMot), Dorsal Attention (DorsAttn), Salient Ventral Attention (SalVentAttn), Limbic, Control/Frontoparietal (Cont), and the Default Mode (Default/DMN) networks. An ROI-ROI connectivity matrix of Fisher-transformed bivariate correlation coefficients between each pair of time series was generated for each participant. The whole-brain matrix contained 400 *choose* 2 = 79,800 connections.

We next examined a selection of well-known meta-analyses on neuroimaging studies in reading and language (in both children and adults) to generate a set of coordinates in MNI space which could then be mapped to ROIs in the Schaefer parcellation. Papers met the following criteria: (1) included only studies performed with structural MRI/fMRI or PET imaging; (2) focused on group contrasts between participants with and without a reading/language impairment, or a functional contrast between reading/language and control tasks; (3) reported results as coordinates in either Talairach or MNI space; and (4) focused on English-speaking participants or cross-cultural results. Talairach coordinates were converted to MNI using the Lancaster transformation implemented in GingerALE (Eickhoff et al., 2009; Laird et al., 2010; Eickhoff et al., 2012). A total of 26 meta-analyses were examined. Of these, 22 reported a total of 416 unique coordinates. These coordinates were mapped to the ROI in the Schaefer template in which that coordinate was spatially contained (referred to as a given coordinate’s ‘parent ROI’). 49/416 mapped onto non-cortical regions, leaving a total of 367 in-range coordinates. These 367 coordinates loaded onto a total of 152 unique Schaefer ROIs (some unique coordinates were mapped onto the same ROI). The Supplementary Material includes a spreadsheet with citation information for the final meta-analyses (Supplementary File 2), and another spreadsheet containing all extracted coordinates and their associated ROIs (Supplementary File 3).

#### 2.5.1 ROI cutoffs

To investigate the specific contribution of reading and language networks to brain-age predictions, four ROI cutoffs were chosen. Cutoffs were based on the total number of coordinates that mapped to a given Schaefer ROI. The greater the number of unique coordinates that map to a given region, the more likely that region is to be part of the core language/reading network and thus relevant to group differences between participants. We created the following sets of ROIs (Figure 2a-d): 400 ROIs (whole-brain/no cutoff); 152 ROIs (≥1 coordinates mapped to region); 78 ROIs (≥2 coordinates); and 27 ROIs (≥4 coordinates). For the subsequent analyses, pairwise ROI-ROI connections were derived only between included ROIs. As the ROI cutoff increases, the data used to train the models becomes more concentrated in reading network regions.

**Figure 2.**
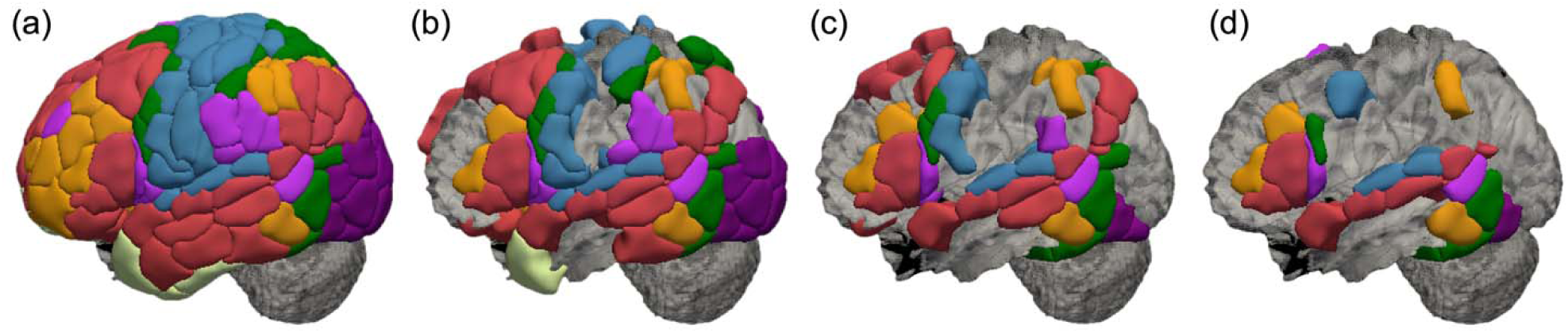
ROI sets generated from literature review, left lateral view. (a) Whole-brain 400-ROI set. (b) 152-ROI set. (c) 78-ROI set. (d) 27-ROI ‘core reading network’ set.

### 2.6. Model training, cross-validation, and testing

Prior to SVR model training, PCA was applied to all FC data in the training set to reduce the number of features while preserving data variance. PCA converts the feature set into a smaller number of uncorrelated principal components (PCs), each a linear combination of correlated original features (Leonardi et al., 2013; Amico & Goñi, 2018). The linear transformation is computed from the top eigenvectors of the covariance matrix, ranking components by explained variance (Khosla et al., 2019). PCA was performed for each set of ROIs using Python’s Scikit-learn, retaining 95% of the original variance.

Each SVR model was built using Scikit-learn. SVR was chosen due to its previous success in constructing age-prediction models from brain data, its resilience to outliers, and its flexibility: it can produce reliable performance even with a relatively small training sample and many features, as is the case for our dataset (Soumya Kumari & Sundarrajan, 2024). A linear kernel was chosen to ease model interpretability, and hyperparameter ε was set to 0.1 such that the model received no penalty in the training loss function for data points predicted within 0.1 years of the true age value. Pre-PCA features were all pairwise ROI-ROI Fisher’s Z-transformed bivariate correlation coefficients between ROIs. The final set of training features consisted only of the PCs which remained after running PCA when preserving 95% of the variance. Reading ability was not used to train the model. Training labels were floating-point numbers describing chronological age for all participants.

For each model, 1000 training permutations were done, each with a unique split for training/cross-validation and hold-out testing data. For each permutation data was split into the training/cross-validation and test sets using a 2-1 train-test ratio (N = 496/N = 246) prior to any model training or cross-validation. Within each permutation, 5-fold cross-validation was done *on the training set only* using Scikit-learn’s built-in method. The splitting procedure was performed for all three groups (PR, TR, ER) separately to ensure that each group was represented proportionally in both the training and testing sets. For each group a random 2-1 train-test split was performed. The three separate training groups were then combined to form a single training group where each group was represented proportionally; the same was done for the testing set. PCA was then performed on the training data only, to ensure that the test set was withheld from all models throughout training and cross-validation. The resulting PCA model was used to transform the training data prior to training/cross-validation, and the same model was used to transform the test set prior to testing.

### 2.7. Analyses

#### 2.7.1. Between-model differences in age prediction (whole-brain vs. reduced-ROI)

We compared prediction accuracy between the whole-brain and reduced-ROI reading network models. Predictions were averaged for each participant across all permutations. All models were compared on their ability to correctly predict age across all participants, regardless of decoding ability, using both traditional error metrics [root mean squared error (RMSE) and mean absolute error (MAE)] and Akaike’s Information Criteria (AIC). AIC estimates prediction error and can be used to compare the predictive ability of non-nested linear statistical models by calculating their relative likelihood. We hypothesized that there would be a positive relationship between model performance (minimized prediction error) and the number of ROI connections (prior to PCA) used to train the model. Therefore, models trained with whole-brain data would more accurately predict the chronological age of the participants, collapsed across all three reading groups.

#### 2.7.2. Identification of the most important ROIs / connections (whole-brain model, exploratory)

For each training/testing permutation, the PCA coefficient matrix (N x M; N = number of PCs, M = number of ROI-ROI connections) and SVR model coefficient array (1 x N) were matrix-multiplied to get an average coefficient for each original ROI-ROI connection (1 x M). These 1 x M arrays were averaged across all permutations to get a final average coefficient for each ROI-ROI connection. The coefficients were then ranked and sorted by absolute value, and the top features which met the threshold for significance (*p* < .05) using permutation testing were selected for further examination. Permutation testing was performed by randomly permuting the coefficient values for all features 2,500 times, then using these 2,500 permutations as a null distribution of coefficients for each feature. The absolute value of a coefficient represents that feature’s importance in model predictions. The top features were then decomposed into their component ROIs, and these ROIs were then pooled together to determine which occurred most frequently in the highest-ranked features. Each ROI’s assigned brain network from the Schaefer parcellation was also extracted. Permutation testing was then used to test whether the most frequent ROIs and networks within the top features were overrepresented compared to random chance; this was done by randomly selecting (with replacement) the same number of ROIs as were present in the set of significant features from the full set of ROIs in the Schaefer parcellation 2,500 times. These randomly-chosen ROIs were also assigned to their respective networks. This gave us 2,500 sets of null ROIs and their networks, each of which should result in a sampling distribution that is representative of the spatial extent of each network. We use these samplings as the null frequency distribution for each ROI and network. For each of the 400 ROIs tested, we used a strict α value of .0001 to negate the impact of Type I error rates and to result in a manageable/interpretable number of the ‘most important’ ROIs.

#### 2.7.3. Between-group differences in the brain-age gap (BAG)

We calculated the BAG for each participant’s average age prediction across all testing permutations, which is quantified as True Age – Predicted Age. Positive BAG values indicate that the model *under*estimates the true age of the subject, while negative values indicate *over*estimation of age. We compared group differences in overall mean BAG and further examined whether this bias interacted with model type (whole-brain vs. reduced-ROI models). We expected that there would be a difference in the BAG based on reading ability (main effect of Group); and that these effects would be preferentially observed in the reduced-ROI models rather than the whole-brain model.

## 3. Results

### 3.1. Between-model differences in age prediction

All raw model predictions were first converted to z-scores, multiplied by the standard deviation (SD) of the sample’s age, and added to the sample’s mean age to ensure predictions were on the same scale as the true age. This was done to ensure that error metrics were not overly inflated while preserving the model’s relative age predictions across participants. Error metrics for all models are shown in Table 2. Both chosen error metrics (RMSE and MAE) were inversely related to the number of ROIs used in feature construction: fewer ROIs used in training resulted in higher model error, as expected.

**Table 2.** Error metrics for each SVR model.

|  | 400-ROI | 152-ROI | 78-ROI | 27-ROI |
| --- | --- | --- | --- | --- |
| RMSE | 2.68 | 2.83 | 2.98 | 3.11 |
| MAE | 2.04 | 2.17 | 2.30 | 2.47 |

AIC results showed that the whole-brain model minimized prediction error for chronological age (Table 3). Best-fit regression lines for all models and corresponding *R^2^* are shown in Figure 3. Thus, we can be confident that the whole-brain model captures a meaningfully larger amount of age-related variance in the functional data compared to the reduced-ROI models, as expected.

**Figure 3.**
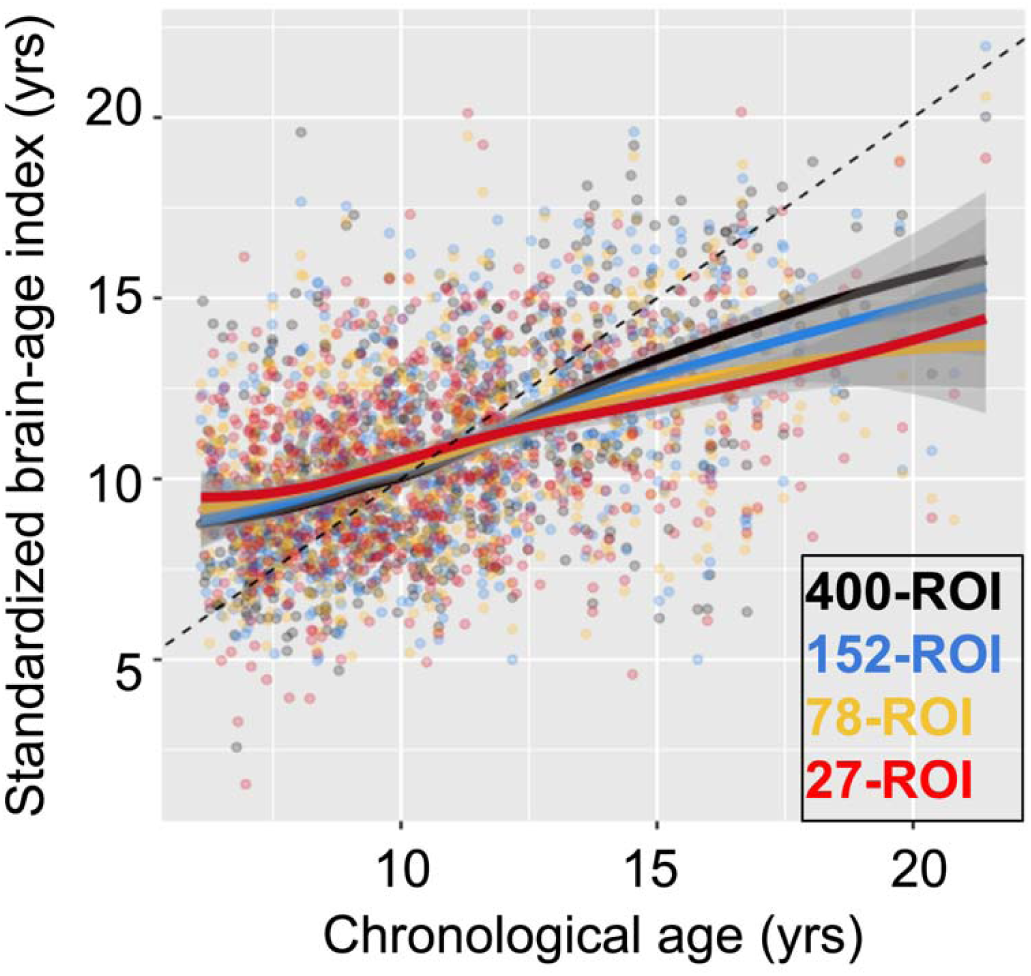
True age (x-axis) vs. standardized brain-age estimate (y-axis) for all models. Dotted reference line is at *y = x*. Lines of best fit for individual models are smoothed.

**Table 3.**
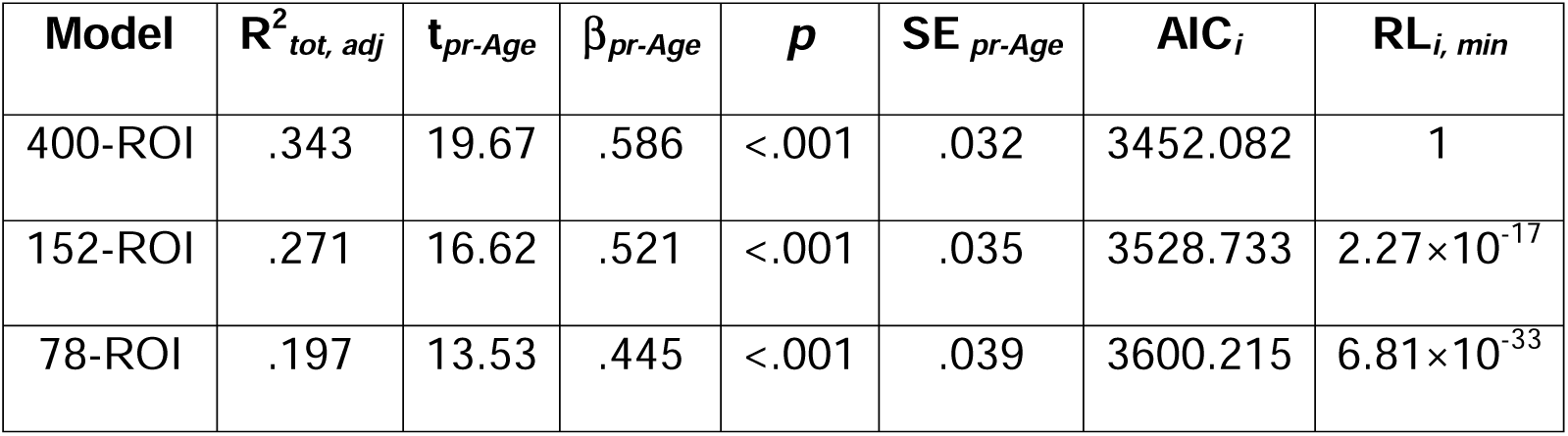
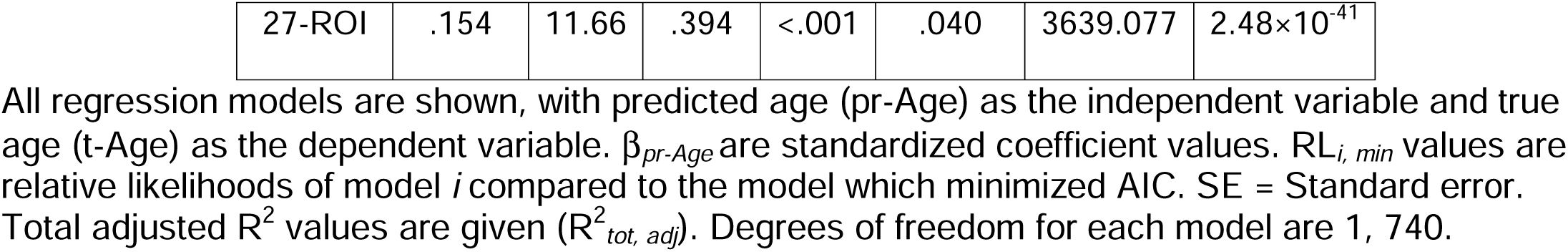
Prediction accuracy for all models (whole-brain and reduced-ROI).

### 3.2. Identification of the most important ROIs / connections (whole-brain model)

#### 3.2.1. Connectivity feature importance

Results from permutation testing showed that only 4,000 features (5.01% of the total feature set) met our criteria for significance in the whole-brain model. Most were between-network (65.9%) rather than within-network (34.1%) (Figure 4a). A plurality of significant connections was interhemispheric (43.8%), followed by left-hemisphere connections (30.5%) and right-hemisphere connections (25.7%). The interhemispheric connections were predominantly between-network, at a rate of 69.9%. However, comparing these rates to those of the entire whole-brain connectivity matrix shows that the within-network connections were actually *overrepresented* in the set of significant features, while between-network connections were underrepresented, regardless of hemispheric identity (Figure 4b). We also visualized those connections with absolute coefficient values >2.5 SDs above the mean and overlaid them onto an MNI-template brain (Figure 4c, left), along with their component ROIs (Figure 4c, right).

**Figure 4.**
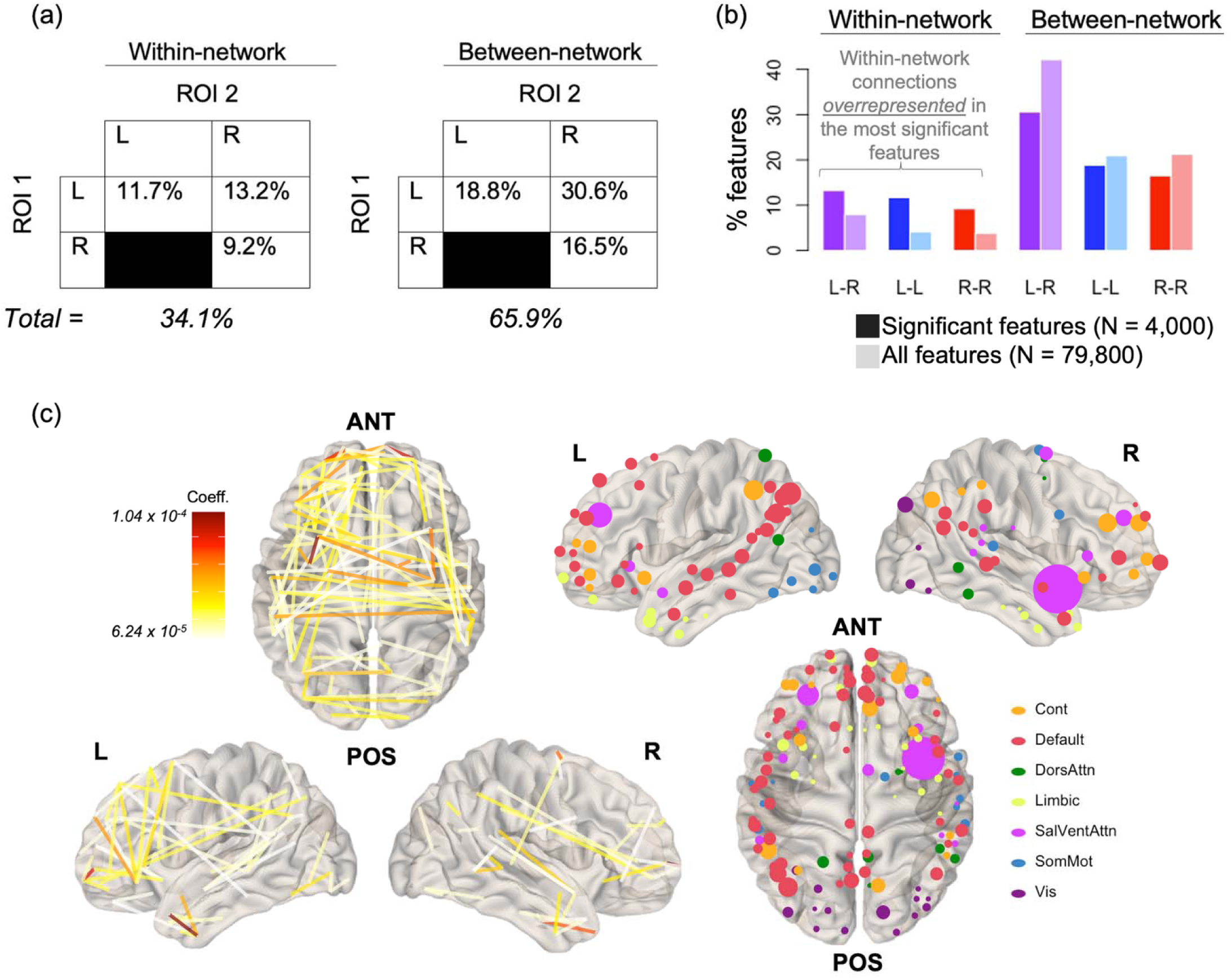
Significant connections and their hemispheric origins. (a) The N = 4,000 significant connections were predominantly between-network (65.9%), interhemispheric (43.8%), or both (30.6%). (b) Comparison of the percentages in (a) to those from the full set of N = 79,800 connections shows that within-network connections are overrepresented among significant features, regardless of hemispheric identity. (c) Connections with coefficient magnitudes >2.5 SDs above the mean (left) and their associated ROIs (right), with ROI size weighted by relative frequency within the significant connections. ANT = Anterior, POS = Posterior.

#### 3.2.2. ROI importance

We found that 36 ROIs in the Schaefer parcellation met our strict permutation-testing criteria for being overrepresented in the set of significant connections (Figure 5a, b). The median frequency of all ROIs in the significant features was 18 (SD = 13.73). Among the 36 significant ROIs, the median frequency within the significant features was 45 (SD = 19.09, range 36-150). These regions were primarily drawn from temporal, frontal, and parietal regions of cortex, with little representation from occipital regions. The set of features associated only with these ROIs (36 choose 2 = 630 connections) had a larger average absolute feature coefficient than the full set of features (Figure 5c). The region with the single greatest frequency of occurrence within the significant features was the frontal operculum/insula (SalVentAttn), which occurred in 3.75% of significant features.

**Figure 5.**
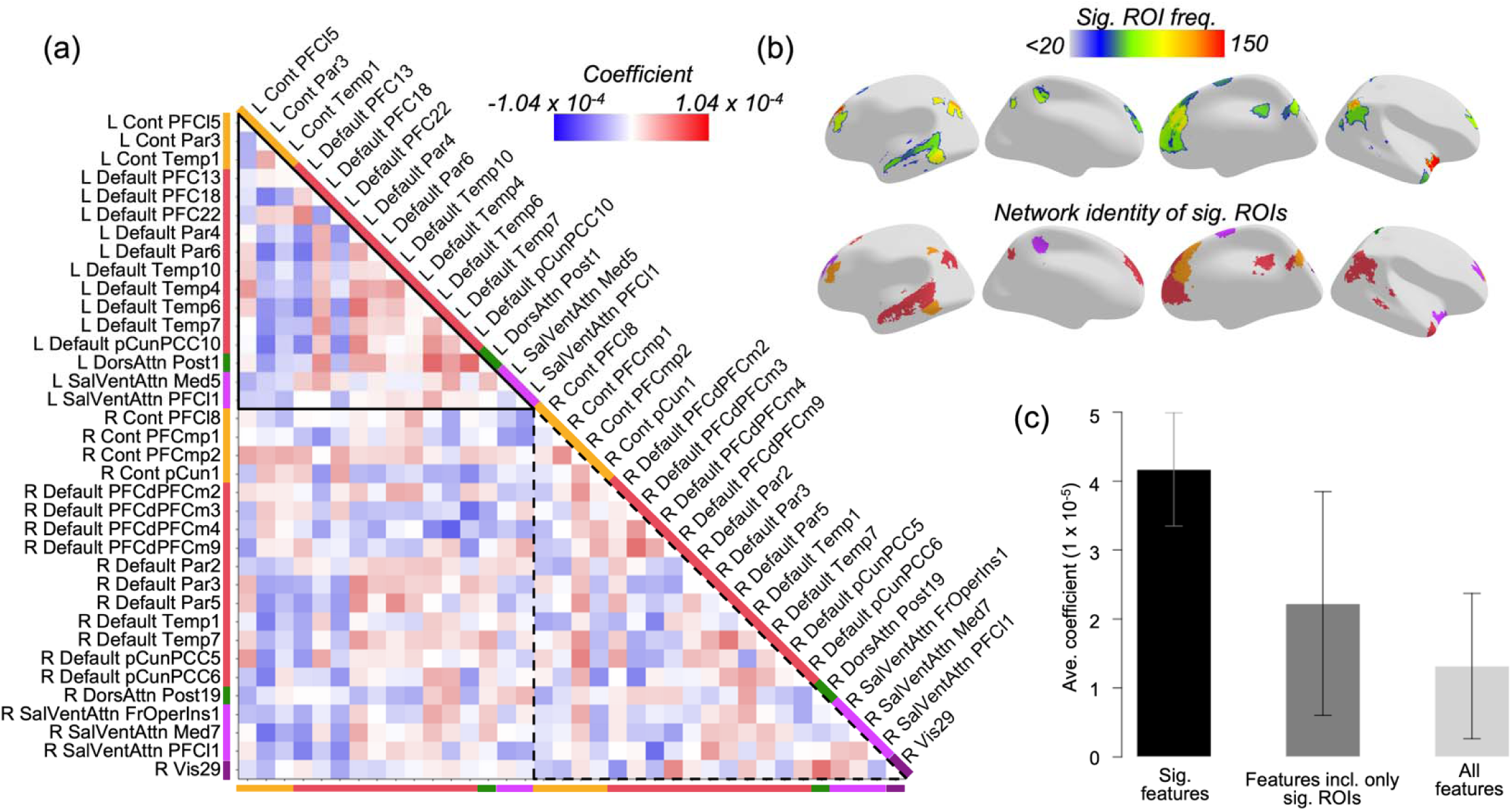
Top ROIs in the whole-brain model. (a) Feature coefficients for each pair of significant ROIs. Colored bars on each side of the triangle reflect each ROI’s assigned network in the Schaefer parcellation. (b) Top row: color represents the frequency of occurrence for each significant ROI in the 4,000 significant connections. Bottom row: color represents the assigned network for each significant ROI. (c) Features including only significant ROIs (middle bar) have higher average coefficient values than all features together (right), but lower average coefficient values than the 4,000 significant features (left).

#### 3.2.3. Network representation

We used the same set of permutations generated for our ROI testing procedure to determine if the significant ROIs/connections came predominantly from specific brain networks. We used the ROI-based permutations due to the fact that certain networks cover a larger spatial area and consist of greater numbers of individual ROIs than others, meaning that their representation in a randomly-selected sample of ROIs will already be skewed.

Results showed that three networks were overrepresented in the significant features (the SalVentAttn, Control, and Default networks), while the other four were underrepresented (the Visual, SomMot, DorsAttn, and Limbic networks) (Figure 6a). Both positive and negative coefficients were dominated by connections involving the DMN, although the nature of those connections differed: the positive connections were dominated by within-network connections of the DMN and other networks, while negative ones were dominated by connectivity between the DMN and the Control/SalVentAttn networks (Figure 6b).

**Figure 6.**
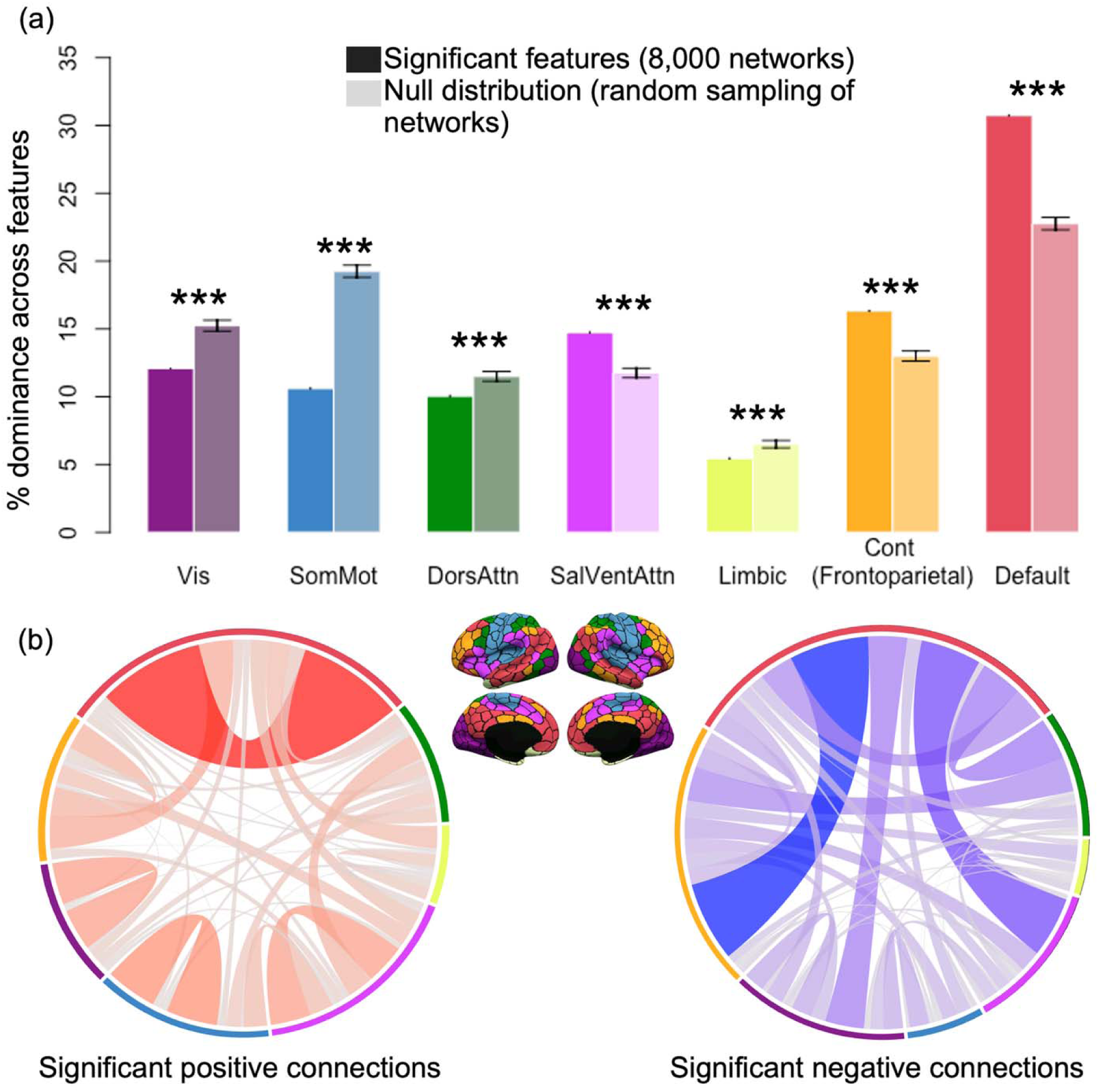
Network representation among the most significant features in the whole-brain model. (a) Darker bars represent a network’s relative dominance in the set of 8,000 ROIs derived from the 4,000 significant connections. Lighter bars represent a network’s average relative dominance across all 2,500 permutations for randomly-chosen ROIs. Error bars are given in SD. ***p < .001. (b) Connectivity patterns differed based on the sign of coefficients.

### 3.3. Between-group differences in the BAG

We ran a repeated measures analysis of covariance (RM-ANCOVA) on BAG values, with Model (400-ROI, 152-ROI, 78-ROI, 27-ROI) as a within-subjects effect, Group (ER vs. TR vs. PR) as a between-subjects effect, and true age as a covariate of no interest, with interaction effects included. There was a significant effect of Group, F(2,736) = 14.46, *p* < .001, partial η^2^ = .038. However, there was no Model x Group interaction (*p* = .483), implying that the group effect on BAG was not clearly different between models. Post-hoc ANCOVA follow-ups in each model separately revealed that the effect of Group was significant across all models: 400-ROI: F(2,736) = 14.74, *p* < .001, partial η^2^ = .039; 152-ROI: F(2,736) = 12.89, *p* < .001, partial η^2^ = .034; 78-ROI: F(2,736) = 10.22, *p* < .001, partial η^2^ = .027; 27-ROI model: F(2,736) = 6.76, p = .001, partial η^2^ = .018. Contrary to our hypothesis, the 400-ROI whole-brain model had the most robust effect of Group. These effects were driven primarily by a more negative BAG in the ERs (overestimation of age) and a more positive BAG in the PRs (underestimation of age) (Figure 7). Therefore, advanced readers’ ages were more likely to be overestimated by the model trained only on reading/language network ROIs, while impaired readers’ ages were more likely to be underestimated, as expected.

**Figure 7.**
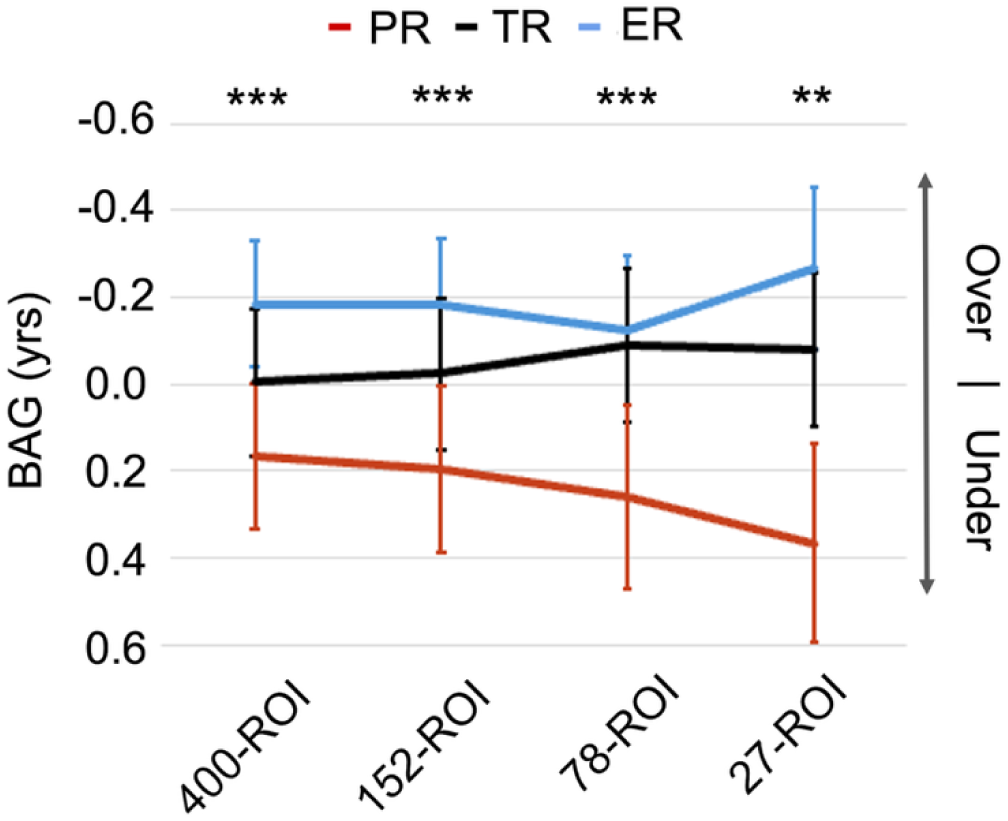
Mean brain-age gap values for all models, separated by reading group. Positive values indicate underestimation of true age, while negative values indicate overestimation. Line color indicates reading group. Error bars are given in standard error. Significance indicates a Group effect on BAG. ***p < .001, **p < .01.

## 4. Discussion

We successfully trained a model to predict age based on FC data and showed bias effects in model predictions based on reading status. Connections involving frontoparietal control regions and the DMN were the most heavily weighted in predicting age. Analyses confirmed the hypotheses that (1) the whole-brain model is a better predictor of age overall than the reduced-ROI models, and (2) there is a group bias in the model such that the ages of PRs are underestimated, and the ages of ERs are overestimated, when controlling for true age. However, in contrast to our expectation that this group bias would be strongest in the 27-ROI model, the opposite trend was observed: the whole-brain model showed the greatest group effects on the brain-age gap.

The exploratory results suggest that regions and connections with the most predictive value come from the DMN and the frontoparietal control networks. Interconnectivity within regions in the DMN are known to strengthen across childhood development (Buckner et al., 2008; Fair et al., 2008) and reflect advancement in a variety of cognitive processes, including reading (Bailey et al., 2018). While the frontoparietal control network is thought to index cognitive control and executive functioning – both key areas of cognitive development from childhood to adulthood – a role for frontoparietal regions has long been posited in reading, particularly RD (Hoeft et al., 2006, 2010; Schwarze et al., 2023; Mascheretti et al., 2024). A slim majority of the densest reading-related regions from our literature review in this study were identified as belonging to frontoparietal or cingulo-opercular control networks, and the DMN (14/27 ROIs). Taken together, the overlap between regions most implicated in reading meta-analyses and those identified as contributing the most to age predictions suggests a strong link between reading maturity and age-related brain development (Vogel et al., 2013). Regarding how these networks map onto reading processes, the frontoparietal network is a key component of the dorsal visual stream, and disturbances in connectivity could interfere with both attentional and perceptual processes, which have long been thought to be atypical in RD (Mascheretti et al., 2024). The DMN contains regions long implicated in reading-related activation, such as the temporal lobes and angular gyrus (Martin et al., 2015; Bailey et al., 2018); both of which overlapped with significant ROIs in the whole-brain model. It seems, then, that the brain networks subserving reading undergo major development in their connectivity patterns from early childhood to late adolescence, resulting in their relative dominance in predicting age in this sample; as such, it is likely that this degree of overlap may not be present when studying development from late adolescence into adulthood (when reading development plateaus).

This period of childhood development is characterized by relative strengthening of co-activation between regions within the same functional network (particularly in the DMN and frontoparietal control networks). Our results also confirm prior findings that show anticorrelated connectivity between the control network and DMN from childhood to late adolescence, given that our most negative significant connections were disproportionately connections between the control network and the DMN (Figure 6b; see also DeSerisy et al., 2021). These results are reflective of these networks’ status as ‘task-active’ vs. ‘task-inactive’, respectively. Altogether, we have observed a relative strengthening of local connections throughout early childhood and adolescence, with a disproportionate emphasis on connectivity within functional networks, although between-network connectivity still plays a dominant role during development in this stage of life.

The whole-brain model was a better overall predictor of age than the other three reduced-ROI models. Its predictions accounted for more variance in the participants’ true age, and it had a much higher relative likelihood than the other models. It is worth noting that the level of variance explained between the whole-brain model collapsed across groups (R^2^ = .343) and the most reduced-ROI model with 27 ROIs (R^2^ = .155) decreased only by approximately half, despite the models’ features being constructed from vastly different amounts of data. The total number of connections used to construct features for the 27-ROI model (351 pairwise connections) was <.005% of the amount used for the whole-brain model (79,800 connections). Yet, this model was still able to capture a significant amount of variance in age based on connectivity between a highly restricted set of brain regions and still showed a bias effect based on RD status. This suggests that while training with widely-diffused whole-brain data will result in more accurate age estimates, accuracy is not linearly proportional to the total number of ROIs or connections used to generate features. Markers of age, in addition to markers of reading ability, are diffusely represented in connectivity patterns across the brain.

We also predicted that there would be a larger group bias in our model’s age predictions when it was trained to predict age using FC data from reading network ROIs drawn preferentially from left-hemisphere reading and language regions rather than diffuse whole-brain connectivity data. Contrary to our expectations, the effect of Group was actually strongest in the whole-brain model, and examination of each group separately showed the expectation direction of effect: the ages of PRs were underestimated, and the ages of ERs were overestimated, while the TRs had an average BAG closer to 0. However, the PRs and ERs also had a lower absolute BAG and higher R^2^ values than the TRs, meaning that while the average error in these groups was skewed positive or negative the magnitude of the error was lower compared to the TRs. This may be interpreted as the trained models being better fit (but with greater bias) to data from the PRs and ERs or, alternatively, the TR group having a higher amount of variability within the population, which the models struggled to capture (Supplementary File 4). A possible follow-up to these results may be to examine whether the same functional connections identified as driving the model’s predictions in our exploratory analysis also differ between these reading groups. Such an analysis could utilize a more traditional GLM-based statistical approach, using both resting-state and task-based data.

There are several limitations to the current study. First, the SVR model coefficients are agnostic towards the initial sign or magnitude of the correlation between two ROIs. Bearing this in mind, the developmental trajectories of highly-weighted connections are only explored in terms of either increasing or decreasing with age, rather than more precise statements about their initial state and developmental dynamics (which may be nonlinear). Another point of criticism is that we have trained the SVR model to predict age using a representative split from the PR, TR, and ER populations. Therefore, the model was trained to predict the age of poor (and exceptional) readers as much as it was trained to predict the age of controls. An alternative approach that uses the latter method of training only with controls may produce clearer bias effects. Another approach may also involve matching the groups more strictly on various criteria, including demographic variables.

Furthermore, we do not assess the model’s ability to predict reading skills directly. A follow-up study that instead assesses the model’s ability to predict continuous reading scores or diagnostic status rather than age would be more enlightening about which functional networks in the brain underlie reading ability uniquely. Age would likely still need to be included as a control variable or used as inclusion criteria, since we have noted that chronological age is associated even with age-standardized TOWRE scores in our sample. An alternative approach would be to split participants into narrower age bands after training and then examine whether there are group differences in the model’s age estimates at these different age bins. This would allow us to not only compare age-matched subgroups but would also be informative as to whether there are any developmental patterns to these differences (e.g., present in early childhood but then diminish later in life).

## 5. Conclusion

This study illustrates that models trained on FC data are able to reliably predict participants’ true age. Both poor and exceptional readers show the expected bias in their age predictions: poor readers’ ages are underestimated, and exceptional readers’ ages are overestimated. Future research could focus on refining this model and/or predicting reading skills as opposed to age in order to identify ‘age-agnostic’ brain networks underlying reading development. Integrating machine learning with neuroimaging provides promising avenues for advancing diagnostic and intervention strategies for children with neurodevelopmental and learning disorders.

## Supporting information

Supplementary File 1. Additional preprocessing details.

Supplementary File 2. Meta-analyses used to extract reading-related regions.

Supplementary File 3. Coordinate data, regional information, and metadata for all extracted coordinates from meta-analyses.

Supplementary Results.

---

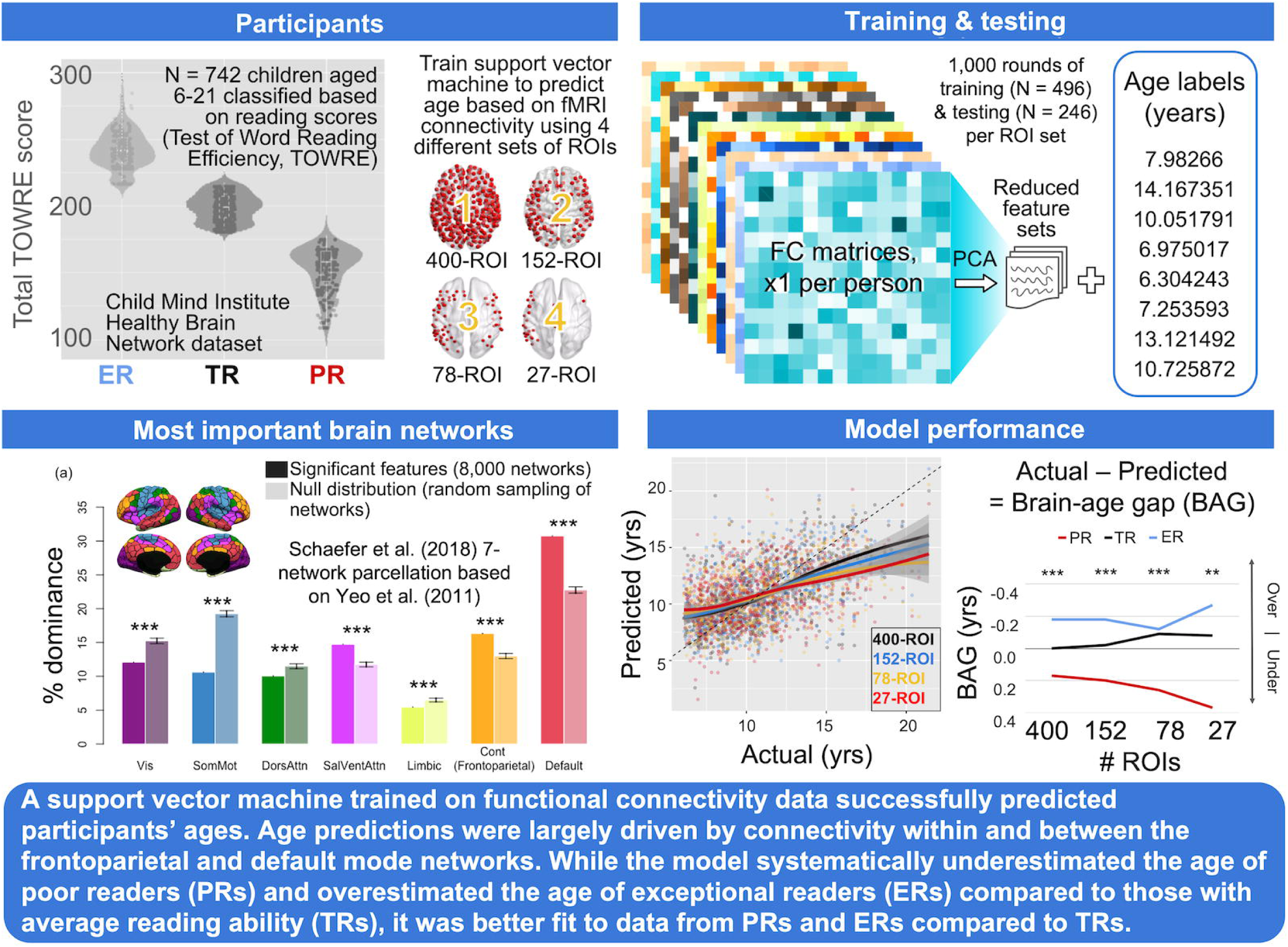

