## Supplementary File 1. Additional preprocessing details. for "Modeling differences in neurodevelopmental maturity of the reading network using support vector regression on functional connectivity data"

**Additional Scan Acquisition & Preprocessing Details**

**Scanner acquisition parameters**

Data (T1-weighted [T1w] structural scans and functional data) was collected at three possible scan locations. Acquisition parameters at each site were as follows:

(1)	Staten Island (SI). 1.5T Siemens Avanto scanner (T1w: repetition time [RT] = 2730 ms, echo time [ET] = 1.64 ms, flip angle = 7°, 176 slices, 1.0×1.0×1.0mm voxels; fMRI: RT = 1450 ms, ET = 40 ms, flip angle = 55°, 54 slices, 2.5×2.5×2.5mm voxels).

(2)	Rutgers University Brain Imaging Center (RU). Siemens 3T Tim Trio scanner (T1w: RT = 2500 ms, ET = 3.15 ms, flip angle = 8°, 224 slices, 0.8×0.8×0.8mm voxels; fMRI: RT = 800 ms, ET = 30 ms, flip angle = 31°, 60 slices, 2.4×2.4×2.4mm voxels).

(3)	CitiGroup Cornell Brain Imaging Center (CG). Siemens 3T Prisma scanner (T1w: RT = 2500 ms, ET = 3.15 ms, flip angle = 8°, 224 slices, 0.8×0.8×0.8mm voxels; fMRI: RT = 800 ms, ET = 30 ms, flip angle = 31°, 60 slices, 2.4×2.4×2.4mm voxels).

**Note**: fMRIPrep information was copied from the boilerplate output log after preprocessing was complete.

**Anatomical data preprocessing (fMRIPrep)**

The T1-weighted (T1w) image was corrected for intensity non-uniformity (INU) with N4BiasFieldCorrection (Tustison et al., 2010), distributed with ANTs 2.2.0 (Avants et al., 2008), and used as T1w-reference throughout the workflow. The T1w-reference was then skull-stripped with a Nipype implementation of the antsBrainExtraction.sh workflow (from ANTs), using OASIS30ANTs as target template. Brain tissue segmentation of cerebrospinal fluid (CSF), WM and GM was performed on the brain-extracted T1w using *fast* (FSL 5.0.9, Zhang et al., 2001). Volume-based spatial normalization to one standard space (MNI152NLin6Asym) was performed through nonlinear registration with antsRegistration (ANTs 2.2.0), using brain-extracted versions of both T1w reference and the T1w template. The following template was selected for spatial normalization: ICBM 152 Nonlinear Asymmetrical template version 6 (TemplateFlow ID: MNI152NLin6Asym).

**Functional data preprocessing (fMRIPrep)**

For each blood oxygen level dependent (BOLD) run, the following preprocessing was performed. First, a reference volume and its skull-stripped version were generated using a custom methodology of fMRIPrep. A deformation field to correct for susceptibility distortions was estimated based on two echo-planar imaging (EPI) references with opposing phase-encoding directions, using 3dQwarp (AFNI 20160207). Based on the estimated susceptibility distortion, an unwarped BOLD reference was calculated for a more accurate co-registration with the anatomical reference. The BOLD reference was then co-registered to the T1w reference using FLIRT (FSL 5.0.9) (Jenkinson & Smith, 2001) with the boundary-based registration (Greve & Fischl, 2009) cost-function. Co-registration was configured with 9 degrees of freedom to account for distortions remaining in the BOLD reference. Head-motion parameters with respect to the BOLD reference (transformation matrices, and six corresponding rotation and translation parameters) are estimated before any spatiotemporal filtering using MCFLIRT (FSL 5.0.9) (Jenkinson et al., 2002). BOLD runs were slice-time corrected using 3dTshift from AFNI 20160207 (Cox & Hyde, 1997). The BOLD time-series (including slice-timing correction when applied) were resampled onto their original, native space by applying a single, composite transform to correct for head-motion and susceptibility distortions. These resampled BOLD time-series will be referred to as ‘preprocessed BOLD in original space’, or just ‘preprocessed BOLD’. The BOLD time-series were resampled into standard space, generating a preprocessed BOLD run in MNI152NLin6Asym space. First, a reference volume and its skull-stripped version were generated using a custom methodology of fMRIPrep. Several confounding time-series were calculated based on the preprocessed BOLD: framewise displacement (FD), DVARS, and three region-wise global signals. FD and DVARS are calculated for each functional run, both using their implementations in Nipype following the definitions by Power et al. (2014). The three global signals are extracted within the CSF, the WM, and the whole-brain masks. Additionally, a set of physiological regressors were extracted to allow for component-based noise correction (CompCor, Behzadi et al., 2007). Principal components are estimated after high-pass filtering the preprocessed BOLD time-series (using a discrete cosine filter with 128s cut-off) for the two CompCor variants: temporal (tCompCor) and anatomical (aCompCor). tCompCor components are then calculated from the top 5% variable voxels within a mask covering the subcortical regions. This subcortical mask is obtained by heavily eroding the brain mask, which ensures it does not include cortical GM regions. For aCompCor, components are calculated within the intersection of the aforementioned mask and the union of CSF and WM masks calculated in T1w space, after their projection to the native space of each functional run (using the inverse BOLD-to-T1w transformation). Components are also calculated separately within the WM and CSF masks. For each CompCor decomposition, the *k* components with the largest singular values are retained, such that the retained components’ time series are sufficient to explain 50% of variance across the nuisance mask (CSF, WM, combined, or temporal). The remaining components are dropped from consideration. The head-motion estimates calculated in the correction step were also placed within the corresponding confounds file. The confound time series derived from head motion estimates and global signals were expanded with the inclusion of temporal derivatives and quadratic terms for each (Satterthwaite et al., 2013). Frames that exceeded a threshold of 0.5mm FD or 1.5 standardized DVARS were annotated as motion outliers. All resamplings can be performed with a single interpolation step by composing all the pertinent transformations (i.e., head-motion transform matrices, susceptibility distortion correction when available, and co-registrations to anatomical and output spaces). Gridded (volumetric) resamplings were performed using antsApplyTransforms (ANTs), configured with Lanczos interpolation to minimize the smoothing effects of other kernels (Lanczos, 1964). Non-gridded (surface) resamplings were performed using mri_vol2surf (FreeSurfer).
