## Supplementary Results. for "Modeling differences in neurodevelopmental maturity of the reading network using support vector regression on functional connectivity data"

**Analysis of demographic covariates**

We tested for group differences in the following demographic variables: age, gender, SES, handedness, WISC-V FSIQ, and the WISC-V block design (BD) subtest, which is a metric for PIQ (see Table 1). We tested continuous variables using one-way ANOVA with Bonferroni correction for multiple comparisons. If Levene’s test was significant below the .05 alpha level, we reported Welch’s F-statistic and used Games-Howell correction for multiple comparisons. SES, an ordinal variable coding for annual income brackets, was compared using the nonparametric Kruskal-Wallis test and Mann-Whitney U tests as pairwise follow-ups. We tested categorical variables (gender and handedness) using Chi-squared tests.

We found a small-to-medium group difference in FSIQ (η^2^ = .298), a smaller effect for PIQ (η^2^ = .093), and a small effect for SES (η^2^ = .060), using effect size interpretations outlined in Sawilowsky (2009). Previous findings have shown that (a) both verbal and nonverbal measures of IQ are associated with and predictive of later reading ability, including dyslexia diagnoses (van Bergen et al., 2014); that (b) verbal IQ measures are more strongly related to reading and language deficits compared to nonverbal measures (van Bergen et al., 2014); and that (c) RD is disproportionately more likely to be identified in children from low-SES environments (Piotrowska & Willis, 2019). Our results here provide additional support for these findings and justifies these variables’ inclusion as nuisance regressors in subsequent analyses. We selected PIQ instead of FSIQ as the IQ regressor in the analysis because FSIQ is more likely to be associated with reading/language skills and its inclusion might remove meaningful variance associated with reading skill. Neither handedness nor gender ratios differed between groups and were also included as covariates of no interest in all analyses. There was also a very small effect of age (η^2^ = .015). Post-hoc tests showed that the significant difference in age was driven by the age gap between ERs and PRs (p <.05), with poor readers being older on average in this sample, while the TRs were intermediate in age and not different from the other two in post-hoc follow-ups (both ps >.05). We therefore consider the effect of age in subsequent analyses.

**Follow-up BAG analysis**

As a follow-up to our main BAG analysis, we also compare group differences in absolute BAG magnitudes; this approach allows for the possibility that one group has excessively high BAG in both directions (i.e., generally higher prediction error), which may result in a group mean BAG of 0.

Results from a series of one-way ANCOVAs performed on both positive and negative BAG values separately for all models is shown in Table S1. Both Group and actual age (and their interaction) are used as predictors. Overall, results suggest that TRs have a higher overall model error than the other two groups. This is corroborated by examining the corresponding R^2^ values separately for each group: PRs = .558, .434, .311, .240 for all models; TRs = .118, .084, .059, .047; ERs = .566, .494, .382, .286. Altogether, this suggests that the models are not as well fit to the TR data.

**Table S1. Summary of absolute BAG values (magnitudes) by group and model.**

|  | abs(BAG) | | |
| --- | --- | --- | --- |
|  | PR | TR | ER |
| 400-ROI* | 1.89 (1.52)_a_ | 2.38 (2.04)_b_ | 1.60 (1.24)_a_ |
| 152-ROI** | 2.10 (1.80)_a_ | 2.48 (2.02)_b_ | 1.71 (1.31)_a_ |
| 78-ROI*** | 2.29 (2.06)_a,b_ | 2.53 (2.01)_a_ | 1.90 (1.44)_b_ |
| 27-ROI** | 2.60 (2.07)_a,b_ | 2.60 (1.93)_a_ | 2.09 (1.56)_b_ |

Standard deviation (SD) is given in parentheses. Different subscripts indicate group significant differences in Bonferroni-corrected post-hoc tests at the .05 level. Models marked with an asterisk (*) show a significant effect of Group when controlling for true age and the Group x t-Age interaction, df = 2, 736. Post-hoc pairwise tests do not control for age. ***p < .001, **p < .01, *p < .05.

**Follow-up connectivity analyses**

We observed a modest inverse relationship between a connection’s feature weight and its spatial range (calculated as the three-dimensional distance between the centroids of that feature’s ROIs), Pearson’s r = -.096, p < .001. Connections whose component ROIs were a shorter spatial distance from one another tended to have higher absolute weights. This effect was observed for both between-network (r = -.083, p < .001) and within-network connections (r = -.042, p < .001).
